# Dynamic Laser Speckle Imaging

**DOI:** 10.1101/626515

**Authors:** Dmitry D. Postnov, Jianbo Tang, Sefik Evren Erdener, Kıvılcım Kılıç, David A. Boas

## Abstract

Utilizing a high-speed camera and recording back-scattered laser light at more than 20,000 frames per second, we introduce the first wide-field dynamic laser speckle imaging (DLSI) in which we are able to quantify the laser speckle intensity temporal auto-correlation function *g*_2_(*τ*) for every pixel individually to obtain a quantitative image of the dynamics of the light scattering particles in the sample. The ability to directly and quantitatively measure the intensity auto-correlation function allows us to solve the problem of how to quantitatively interpret data measured by laser speckle contrast imaging (LSCI), multi-exposure laser speckle imaging (MESI) and laser Doppler flowmetry (LDF). The intensity auto-correlation function is related to the field temporal auto-correlation function *g*_1_(*τ*), which has been quantitatively related to the dynamics of the light scattering particles including flowing red blood cells. The form of *g*_1_(*τ*) depends on the amount of light scattering (i.e. single or multiple scattering) and the type of particle motion (i.e. ordered or unordered). Although these forms of the field correlation functions have been established for over 30 years, there is no agreement nor experimental support on what scattering and motion regimes are relevant for the varied biomedical applications. We thus apply DLSI to image cerebral blood flow in mouse through a cranial window and show that the generally accepted form of *g*_1_(*τ*), is applicable only to visible surface vessels of a specific size (20 – 200*μ*m). We demonstrate that for flow in smaller vessels and in parenchymal regions that the proper *g*_1_(*τ*) form corresponds with multiple scattering light and unordered motion which was never considered to be relevant for these techniques. We show that the wrong assumption for the field auto-correlation model results in a severe underestimation of flow changes when measuring blood flow changes during ischemic stroke. Finally, we describe how DLSI can be integrated with other laser speckle methods to guide model selection, or how it can be used by itself as a quantitative blood flow imaging technique.

## Introduction

The development of faster cameras and sensing techniques is enabling various advances in biomedical imaging applications such as SCAPE microscopy for high speed volumetric imaging^1^ and functional ultrasound imaging^2^. Utilizing a high-speed camera and recording back-scattered laser light at more than 20,000 frames per second, we introduce the first wide-field dynamic laser speckle imaging (DLSI) in which we are able to quantify the laser speckle intensity temporal auto-correlation function for every pixel individually to obtain a quantitative image of the dynamics of the light scattering particles in the sample. In biomedical applications, the dynamic light scattering particles are typically moving red blood cells. Until now, it was possible to obtain the full temporal sampling of speckle intensity fluctuations for single pixels (i.e. channels of data) as is done with laser Doppler flowmetry^3–6^ and diffuse correlation spectroscopy^6, 7^ measurements of blood flow. Scanning laser Doppler flowmetry was introduced to obtain images of blood flow^8^, but the temporal and spatial resolution was limited. Improved temporal resolution for images of blood flow was achieved with laser speckle contrast imaging (LSCI)^9–12^ by measuring the spatial speckle contrast reduction due to speckle intensity fluctuations during the camera integration time. LSCI has had broad impact, for instance enabling the connection between migraine aura and headache^13^.

Because LSCI does not measure the full temporal dynamics of the speckle intensity fluctuations, it is not possible to confirm from model fitting of the data that the correct model of light scattering and particle dynamics is being used to quantify blood flow. The question of the appropriate model to use for analyzing LDF and LSCI measurements of blood flow has been debated for decades^12, 14–17^. Multi-exposure speckle image (MESI)^10, 18, 19^ has been introduced to probe the temporal dynamics of the speckles *indirectly* by changing the exposure time to measure the impact on speckle contrast. MESI was first used to quantify the impact of static scattering to improve estimates of relative blood flow changes^10^. It was then further extended to probe Gaussian and Lorentzian models of the field correlation function^18^, but did not show significant difference between two. DLSI, on the other hand, samples the fluctuations of the speckle intensity with sufficient temporal resolution that the correct model form can be *directly* estimated from the data.

We measure the speckle intensity temporal auto-correlation function *g*_2_(*τ*) which is related to the field temporal auto-correlation function *g*_1_(*τ*) that we can theoretically derive based on the dynamics of the scattering particles and the amount of light scattering from the dynamic particles. The relation between *g*_2_(*τ*) and *g*_1_(*τ*) depends on the amount of static scattering present in the sample as well as measurement specific parameters related to source coherence^20, 21^, detector speckle averaging^22^, and detector noise^10, 11^. The field temporal auto-correlation function generally takes the simple form of 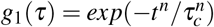 where *τ*_*c*_ is proportional to the time it takes the dynamic light scattering particles to move a wavelength of light, and is thus inversely proportional to the speed of the light scattering particles^9^. The exact form of *g*_1_(*τ*), i.e. the value of *n*, depends on the amount of dynamic scattering, i.e. single or multiple light scattering, and on whether the dynamics are ordered or unordered^15–17^. The amount of dynamic scattering of light for the measurement geometries of LDF and LSCI has typically been assumed to be on average much less than one dynamic scattering event per detected photon^23^, but recent studies have suggested that at least for brain that this number is on average greater than 1^16^. For DCS, the amount of dynamic light scattering is clearly in the multiple scattering regime as the separation between the source and detector on the surface of the tissue is much larger than the scattering length of light within the tissue^6^. For DCS, despite clarity on the amount of dynamic light scattering, the type of motion has been debated with red blood cells expected to have ordered motion but measurements suggesting unordered motion. This discrepancy has only recently been reconciled by a realization that shear induced diffusion of the red blood cells, i.e. unordered motion, is dominating the dynamics of the light scattering particles^7^. It is thus unclear for LDF and LSCI whether the motion of the red blood cells is ordered, as generally expected, or unordered as occurs due to shear induced diffusion^24^.

While the proper *g*_1_(*τ*) model to use for DCS has been established by fitting the actual correlation data^7, 25^, this has never done for LSCI, LDF, and MESI since they do not provide a *direct* measurement of the intensity temporal auto-correlation function. Instead, the *n* = 1 form that corresponds to both multiple scattering ordered motion and single scattering unordered motion has been applied in most LSCI^4, 9, 11^ and MESI^10^ studies. This *n* = 1 form has become the foundation for the simplified relation^26^ between flow speed (*v*) and speckle contrast (*K*), 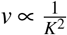, that today is used in the majority of LSCI applications^27–30^ and in the software provided with commercially available LSCI cameras^31–33^. With DLSI we can finally answer the question of what is the correct model to use for *g*_2_(*τ*) and *g*_1_(*τ*) for LSCI and MESI as well as for LDF.

In the present work, we demonstrate DLSI and its application to image cerebral blood flow in the mouse. We show that the generally utilized *n* = 1 form of *g*_1_(*τ*) is applicable only for surface vessels within a specific size range and may lead to a significant misinterpretation of the flow in physiologically relevant applications such as stroke imaging. We discuss other factors that may result in DLSI, LSCI and DCS being more sensitive to unordered motion of red blood cells. Further, while establishing the mathematical foundation for DLSI, we raise concerns regarding the current theoretical framework for analyzing temporal laser speckle contrast imaging results. We then explain how DLSI can be used to calibrate LSCI, converting it to the long sought quantitative tool for imaging blood flow, and how our observations make it possible to improve data interpretation even post-hoc.

## Results

### Intensity autocorrelation

Intensity auto-correlation functions were calculated using Eq.10 and images of *g*_2_(*τ*) for *τ* = 0 and 440*μ*s are shown in Fig.1(C),(D). The *τ* dependence of the correlation functions for four regions of interest are shown in Fig.1(E). For pixels belonging to parenchymal regions, as expected, the correlation function decays slower compared to that from the pixels of surface vessels, which in turn decay slower for the smaller vessels than for the large vessels as the larger vessels generally have faster blood flow. The intensity auto-correlation function at zero time lag, on other hand, varies from location to location, which is not intuitively clear. Instead of it being spatially uniform, as we would initially expect, we see that (i) larger vessels have a reduced *g*_2_(*τ* = 0) value and that (ii) some of the small vessels have *g*_2_(*τ* = 0) values larger than the surrounding parenchyma. To understand this behaviour, we need to explore the theory of speckle decorrelation and the dependence of the dynamics of the scattering particles, on static scattering, and on the effects of spatial and temporal speckle averaging.

**Figure 1.**
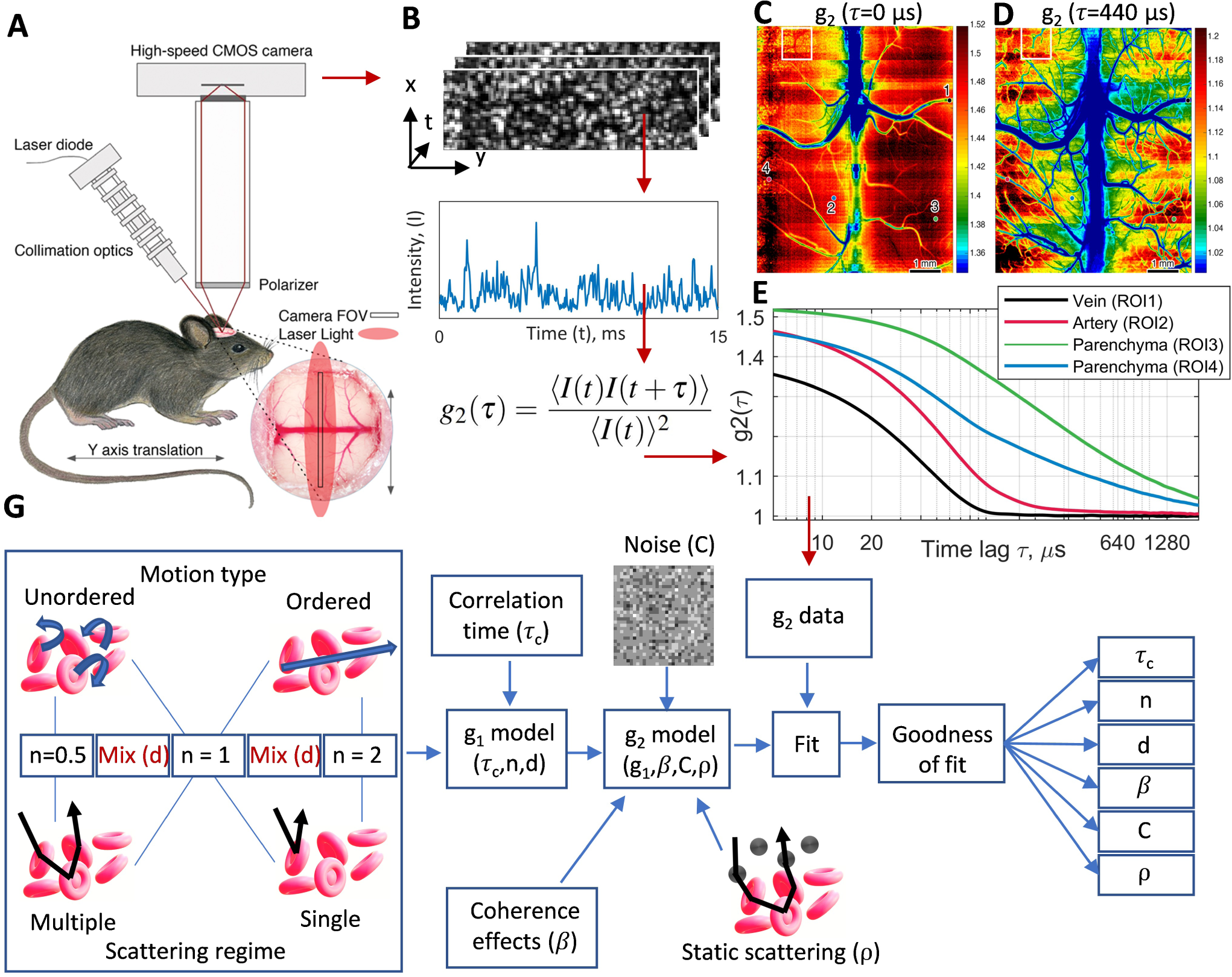
(A) diagram of the imaging system. The collimation optics include a collimator, isolator, anachromatic prism pair, a neutral density filter and beam expander. (B) - data flow (red arrows) and *g*_2_(*τ*) calculation. Panels (C),(D),(E) show the measured intensity temporal autocorrelation function. (C) and (D) show spatial maps of *g*_2_(*τ*) at the specific time lags of *τ* = 0 and 440*μ*s. (E) The *τ* dependence of *g*_2_(*τ*) at selected 3×3 regions of interest that belong to different vessel types as indicated in (C). Notice that not only the decay speed and form is pixel dependent, but also that at *tau* = 0 that some of the vessels appear to have higher values compared to surrounding parenchymal tissue (see the white box in (C)). At later *τ*, the relationship is reversed with lower values of *g*_2_(*τ*) observed in the vessels (see the white box in (D)). (G) Model and fitting process diagram.

### Model

The intensity temporal auto-correlation function *g*_2_(*τ*) is commonly related to the electric field correlation function *g*_1_(*τ*) by the Siegert relation^9, 34^:

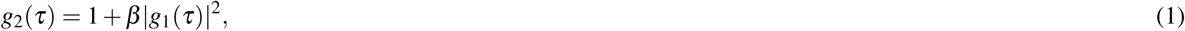

where *τ* is the time lag, and *β* reflects the effects of source coherence and spatial^11, 14, 22^ and temporal averaging on speckle contrast. When measuring blood flow, *g*_1_(*τ*) is generally described^9^ by the form

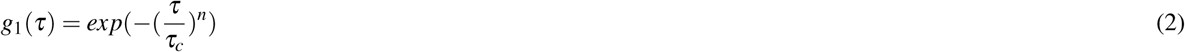

where *τ*_*c*_ is the decorrelation time constant and *n* varies depending on the dynamic light scattering regime^12, 14–16^ (i.e. single versus diffusive light scattering) and the type of motion for the light scattering particles (i.e. ordered versus unordered motion) and takes values of 0.5, 1 or 2. Typical field correlation functions and corresponding scattering and motion regimes are shown in Table1. Model diagram is shown at Fig.1(G)

**Table 1.** Forms of the electrical field temporal autocorrelation function depending on the scattering regime and motion type.

| $g_1$ form | Motion type | Scattering regime | Notation |
| --- | --- | --- | --- |
| $\exp(-(\tau/\tau_c)^2)$ | Ordered | Single | $SO_{n=2}$ |
| $\exp(-\tau/\tau_c)$ | Ordered | Multiple | $MO_{n=1}$ |
| $\exp(-\tau/\tau_c)$ | Unordered | Single | $SU_{n=1}$ |
| $\exp(-(\tau/\tau_c)^{0.5})$ | Unordered | Multiple | $MU_{n=0.5}$ |

The parameter *β* is typically assumed to be homogeneous across the field of view. In DLSI, because of the finite exposure time of the camera *T*_*exposure*_, temporal speckle averaging effects may appear. Specifically, one can expect that *β* will decrease when the exposure time is not much much shorter than the decorrelation time constant *τ*_*c*_. While we do not address this question in detail here, to avoid further confusion we redefine the coherence parameter *β* as

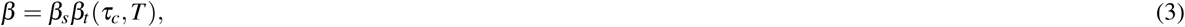

where *β*_*s*_ captures the decrease in *g*_2_(*τ*) arising from spatial speckle averaging^22^ and the partial coherence of the source^20, 21^ and *β*_*t*_ captures effects arising from temporal averaging. While *β*_*s*_ is expected to be constant across the field of view, *β*_*t*_ will depend on sample dynamics at each spatial location and will decrease when the sample dynamics are fast enough such that *tau*_*c*_ is no longer much much longer than the camera exposure time. We expect *β*_*t*_ to decrease for larger vessels where the correlation time is comparable with the exposure time as can be seen by the *g*_2_(0) values in Fig.1 (C). For smaller vessels and parenchymal regions, the *β* values are expected to be more similar.

We fit the data from Fig.1 with the three different dynamic forms of the basic model (i.e. n=0.5, 1, and 2). The *R*^2^ goodness of fit metric for the n=0.5, 1 and 2 fits are shown in Fig.2(D),(E), and (F) respectively. It is evident that different regions are fit better with different dynamic forms for *g*_1_(*τ*). Fig.2(A) shows which form provides the best model fit for each location. We see that the parenchymal regions and smallest vessels are best fit with the model that corresponds with multiple scattering and unordered motion (MU_*n*=0.5_); that the visible surface vessels are best fit with the model of single scattering with unordered motion or multiple scattering with ordered motion (MO/SU_*n*=1_); and that the largest vessels are best fit with the single scattering and ordered motion (SO_*n*=2_) model. The decorrelation time constant *τ*_*c*_ recovered based on the best form fit varies between 50 and 1500 *μ*s, with 50-200 *μ*s appearing for the large vessels, 200-400 *μ*s for the smaller vessels and 400-1500 *μ*s for the parenchymal regions. Contrary to expectations, the *β* fit shown in Fig.2(C), appears to be strongly heterogeneous. While large vessels have the smallest *β* as expected because of the effect of temporal speckle averaging, the larger *β* values for the small vessels compared with the parenchymal regions cannot be explained in terms of temporal or spatial speckle averaging or in terms of source coherence properties. A likely explanation for this behavior is the contribution of static scattering which is likely present in parenchymal regions but is neglected in our simple model fit of the data. Note that the larger *β* values for the smaller vessels than the surrounding parenchyma is consistent with the *g*_2_(*τ* = 0) maps shown in Fig.1(C). Another interesting observation is that by analyzing the intensity auto-correlation function we are able to visualize more vessels compared to that revealed by temporal or spatial laser speckle contrast analysis. The impression appears from the *β* Fig.1(C) and goodness of fit metrics Fig.1(D),(E),(F) which reveal more small vessels than typically seen in speckle contrast images, and proves to be true upon close up inspection of the speckle analysis results (see Fig.11). The reason for this is that model fitting is sensitive to subtle variations in the correlation function form, which may not lead to obvious differences in the first order statistics quantified by speckle contrast analysis. Fig.2(G),(H) show example fits to indicate that some locations fit better than other locations. We expect that the reduced quality of the model fit arises from three major factors not considered in the simple model: (i) measurement noise and insufficient temporal sampling, (ii) static scattering, and (iii) a combination of different sample dynamics.

**Figure 2.**
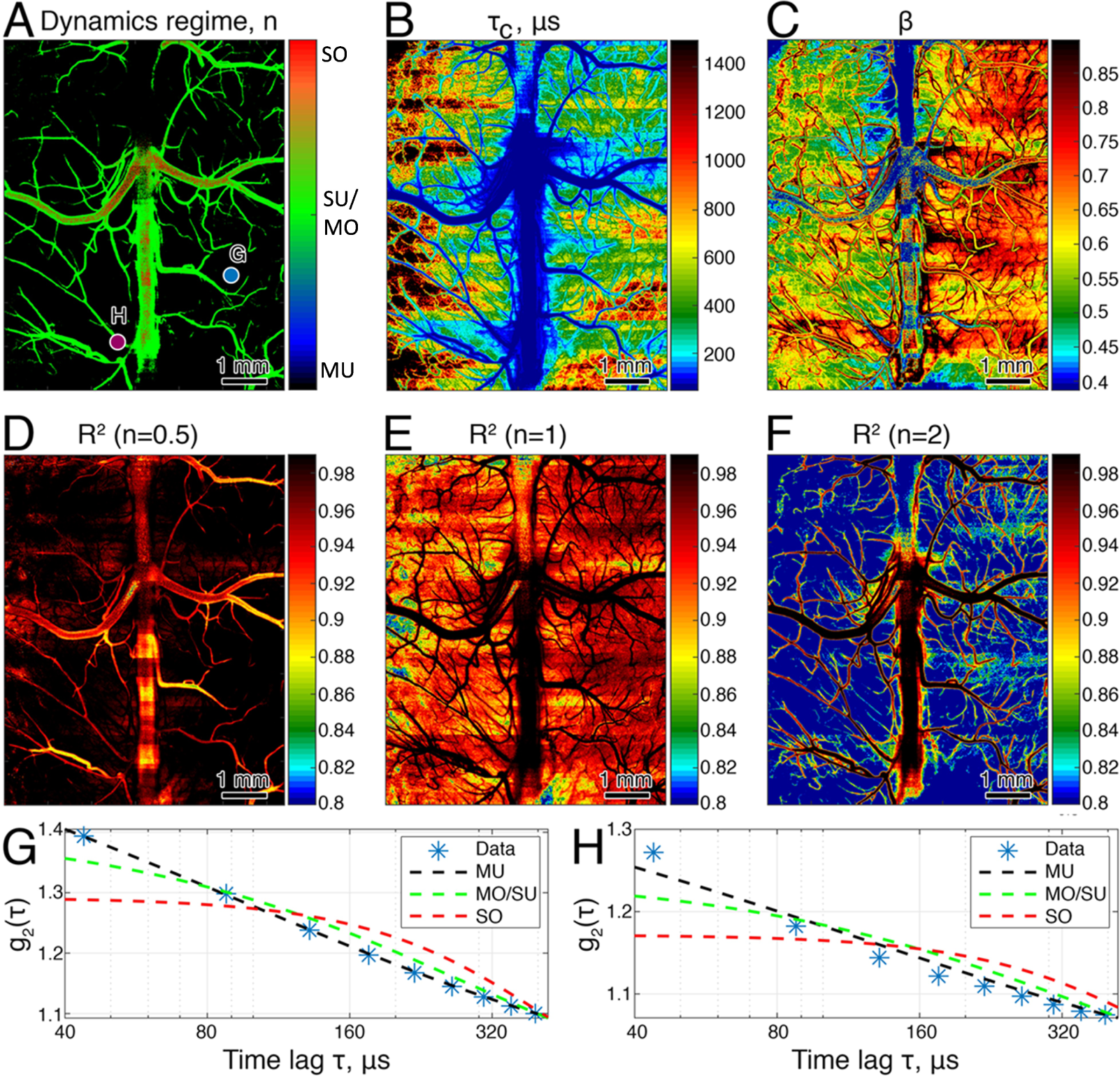
Results of fitting the basic model to the data (Eq.1-2). (A) The dynamic light scattering field auto-correlation form (i.e. *n*) that results in the best fit of the data. (B) and (C) The best fit decorrelation time constant *τ*_*c*_ and coherence parameter *β* respectively. Panels (D), (E), and (F) show the *R*^2^ values corresponding to the different forms of *g*_1_(*τ*) *n* = 0.5, 1, and 2 respectively. It is evident that the parenchyma generally exhibits MU_*n*=0.5_ behaviour, that small and large vessels are fit better with MO/SU_*n*=1_ and SO_*n*=2_ forms respectively. Panels (G) and (H) show fits of *g*_2_(*τ*) for the pixels marked in (A). While the fit in (G) looks almost perfect, it is evident that dynamics in the correlation function shown in (H) cannot be accurately fit with the basic model.

#### Measurement noise and insufficient temporal sampling

The experimentally measured *g*_2_(*τ*) does not always decay to 1 at long lag times, but can decay to values larger and smaller than 1 as we observe in supplemental Fig.8. When measuring spatial speckle contrast, it is known that the presence of measurement noise and static scattering will result in a constant positive offset term^10, 11, 35^. However, when measuring *g*_2_(*τ*) via a temporal intensity auto-correlation measurement, it is known that *g*_2_(*τ*) decays to 1 even in the presence of static scattering unless spatial ensemble averaging is also performed^35–37^. Thus, our observation of*g*_2_(*τ*) not decaying to 1 needs further explanation. The biggest factor is the effect of temporal sampling. With sufficient temporal sampling of the speckle fluctuations, the intensity variance will equal the mean intensity squared, that is < *I*(*t*)*I*(*t* + *τ*) >=< *I*(*t*) >^2^. This condition of sufficient temporal sampling is met when the camera integration time *T*_*exposure*_ is much shorter than the decorrelation time of the speckle intensity *τ*_*c*_, and when the length of sampling *T*_*total*_ is much longer than *τ*_*c*_. As we see in supplemental Fig.8, the distribution of *g*_2_(*τ*) at large values of *τ* becomes more narrowly distributed around a value of 1 when *T*_*total*_ is increased because we are better meeting the condition of sufficient temporal sampling. However, as *T*_*total*_ increases, a small positive bias remains in the distribution. This positive bias is a result of measurement noise (specifically the camera readout noise and dark count) reducing the probability of measuring speckles with zero and very small intensities which skews the temporal intensity distribution such that < *I*(*t*)*I*(*t* + *τ*) > is greater than < *I*(*t*) >^2^ even at long *τ* and long *T*. To capture these effects of measurement noise and insufficient temporal sampling in our model, we consider a form for *g*_2_(*τ*) with a constant offset term *C*

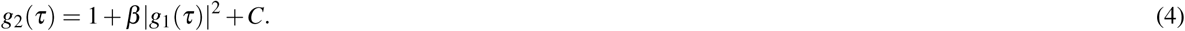

Results of fitting the model with the constant term are shown in Fig.3(A),(B),(C). From the F-test^38^ results comparing the fit with constant offset model (Eq.4) versus the fit with the basic model (Eq.1) as shown in Fig.3(C), it is clear that the model with the constant term *C* performs better than the basic model in almost all regions, except the largest vessels. The spatial map of the best fit for *n* (Fig.3(A)) is similar to the one obtained with the basic model fit. The fit parameter *C* has a higher value in regions with a slower flow (Fig.3(B)). We might expect the regions with slower flow to have a wider distribution in the values of *C* as we see in Fig.8(A),(B),(C) when *T* = 21ms is short and there is less temporal sampling of the slower speckle fluctuations than for the regions with faster speckle fluctuations. But for the experimental results presented here, *T* = 4000ms and Fig.8(G),(H),(I) indicates that there is sufficient temporal averaging for the regions with the slowest flow (Fig.8(G)) as well as for the regions with the fastest flow (Fig.8(I)). Fig.8(G),(H),(I) does indicate that the regions with the slower speckle fluctuations have a larger positive skew in the distribution of values for *C*. Investigation of the temporal speckle intensity distribution for the regions with fast versus slow speckle intensity fluctuations indicates that this difference in the positive skew is a result of the time scale of the intensity fluctuations compared with the camera frame exposure time *T*_*exposure*_ (see supplementary Fig.9). For the faster speckle intensity fluctuations, the camera exposure time smooths the speckle contrast, reducing *beta* in the process and having the effect of reducing our sensitivity to the camera noise resulting in a smaller value for *C*.

**Figure 3.**
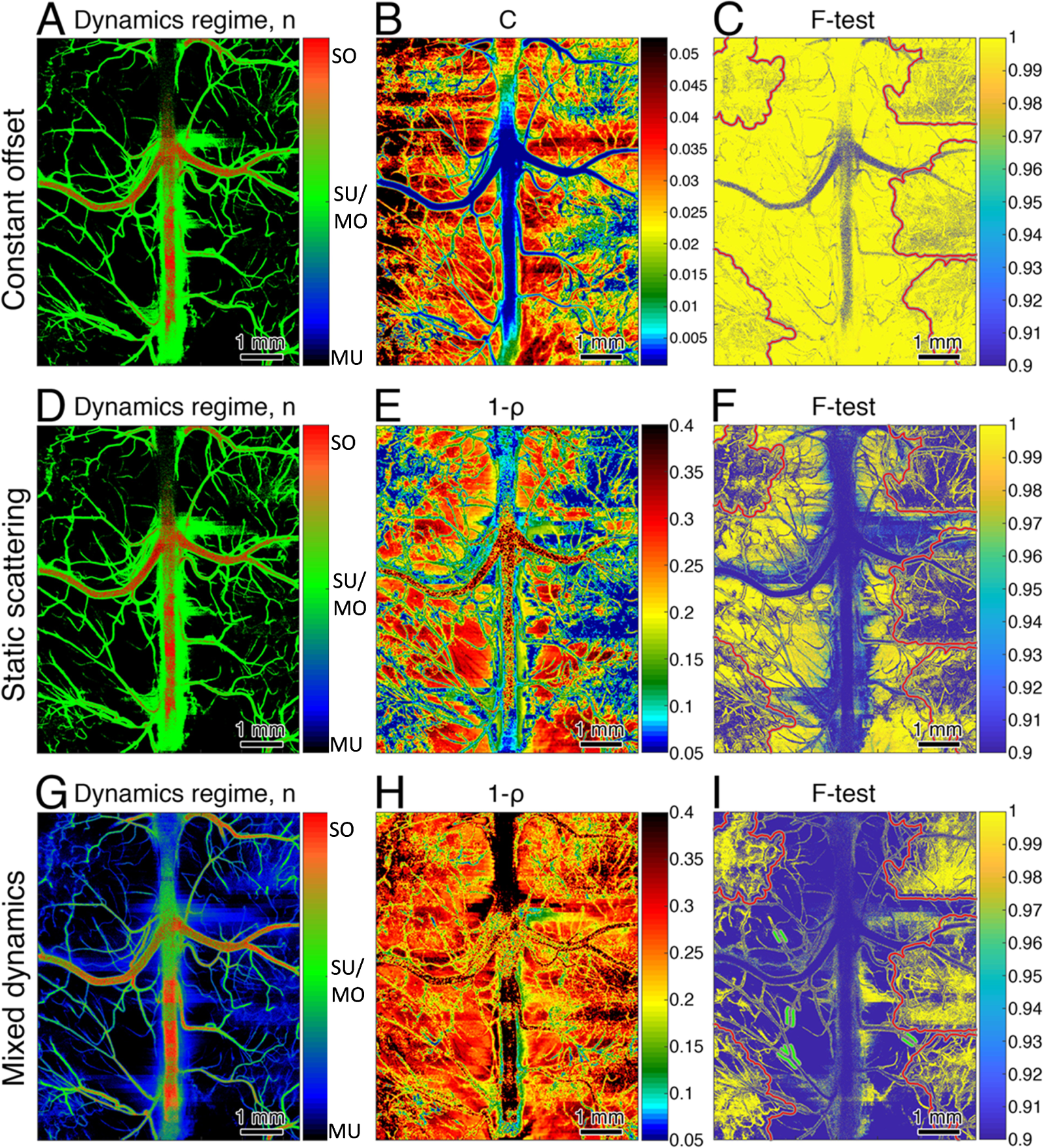
Comparing different model fits. Panels (A), (B), and (C) present results for the model considering the constant offset from measurement noise. (A) shows the best fit for *n*, (B) shows the constant *C* obtained with the best *n*, (C) shows the results of the F-test comparison, shown as F=1-p where p indicates the statistical significance such that the new model is better when F>0.95. Panels (D), (E), and (F) present the results for the model considering static scattering. (E) shows the static scattering component (1 – *ρ*) and (F) shows the results of the F-test comparison with the best fit of the two simpler models. Panels (G), (H), and (I) present results for the mixed dynamics model. Unlike the other models, the form of *g*_1_(*τ*) can take intermediate values between MU_*n*=0.5_ and MO/SU_*n*=1_ or MO/SU_*n*=1_ and SO_*n*=2_, behaviors which are defined by the *d*_*MU*_ and *d*_*SO*_ parameters. Similar to (E), (H) shows the estimated static scattering component. (I) shows the F-test comparison with the best of all other models. Note that the static scattering images (E) and (H) are spatially filtered to reduce fluctuations caused by *I*_*c*_ *≠* ⟨ *I*_*c*_ ⟩. One can see that while (A) and (D) are not qualitatively different from Fig.2(A) (except for more vessels displaying the MO/SU behavior), that (G) instead highlights the presence of transition areas which display both MU and MO/SU behaviors. The transition areas are aligned with the regions where the mixed model proved to be better compared to other models. The same regions were not fit properly by the static scattering model, resulting in unexpectedly reduced static scattering.

#### Static scattering

As we saw above, in Fig.1(C) and (D), some vessels appear to have higher values of*g*_2_(*τ*) at *τ* = 0 that then decay to lower values at long time lags compared to the surrounding tissue. This behavior is the result of a stronger contribution of static scattering in the parenchyma. While both *β*_*s*_ and *β*_*t*_ in the parenchyma should be higher or the same compared to vessels (note that *β*_*t*_ can be reduced in vessels if *T*_*exposure*_ is not significantly shorter than *τ*_*c*_), the contribution of static scattering is greater in the parenchyma, because of a lower blood volume fraction, leading to a decrease in *g*_2_(0). At longer time lags, however, *g*_2_(*τ*) in the parenchyma is higher compared to vessels since the speckle dynamics are slower. Following the theoretical description in^35, 37^, we used a model for *g*_2_(*τ*) that includes the effects of static scattering,

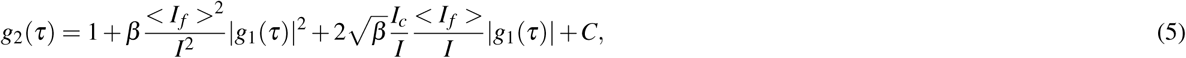

where < *I*_*f*_ > is the temporally averaged intensity of the fluctuating component of the speckle intensity, *I*_*c*_ is the constant component of the static speckle intensity, and *I*_*total*_ =< *I*_*f*_ > +*I*_*c*_ is the total intensity. We denote the fraction of the fluctuating component as 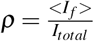. Note that when analyzing the temporal correlation function of the speckle intensity that static scattering does not produce the expected constant offset in the long lag time value of *g*_2_(*τ*). As clearly described in^36, 37^, this is because the temporal analysis does not have access to the proper intensity statistics of the static speckle intensity *I*_*c*_ which can only be accessed by analyzing the spatial statistics. A further consequence of this is that *I*_*c*_ and *I*_*total*_ are not the true ensemble averaged values of the static and total intensity, and their values will vary across space from speckle to speckle. This variation is evident in the total intensity image shown in supplementary Fig.10(C) that despite being an average over 4 sec still reveals a static speckle pattern. As such, the proper average total intensity can only be determined from a spatial average of the local speckle pattern. This has an important consequence for temporal laser speckle contrast imaging (tLSCI) that has not been described in the literature. While tLSCI provides an accurate measurement of the standard deviation of the temporal speckle fluctuations (*σ*), it does not provide an accurate measure of the average total intensity and thus the speckle contrast *K*_*temporal*_ = *σ/I*_*total*_ will not be accurate and will vary unrealistically on the local spatial scale (see supplemental Fig.10(B)). One will have to perform a spatial average of the local speckle intensities to get the proper value for *I*_*total*_ to use in the calculation of *K*_*temporal*_.

Results of using the static model (Eq.5) to fit the experimental data are shown in Fig.3(D),(E),(F). The image of *n* (Fig.3(D)) is comparable to the one obtained using the constant offset model (Fig.3(A)) and using the basic model (Fig.2(A)). One difference that repeats as we increase our model complexity is that more small vessels appear in the images. As shown by the F-test (Fig.3(F)), the static scattering model is significantly better for most of the parenchymal regions compared with the constant offset and basic models, but does not show improvement for large and medium sized surface vessels. This is expected as little static scattering is expected to contribute for measurements from these surface vessels. The amount of static scattering (i.e. 1 – *ρ*) measured from the parenchymal regions varies around 0.2 to 0.3 (Fig.3(E)), which corresponds well with prior results from Multi-Exposure Speckle Contrast Imaging^18, 19^. We do observe some parenchymal regions for which the static scattering model was not better than the constant offset or basic models (regions indicated by red ovals in Fig.3(F)). We expected that a more complex model of the dynamics is needed to explain the data in these regions.

#### Considering mixed dynamics in the form for *g*_1_(*τ*)

Observing the small vessels which display MU_*n*=0.5_ behaviour in the basic model and MO/SU_*n*=1_ behaviour in more complicated models it is reasonable to expect that the transition between these behaviors is not necessarily sharp. Indeed, it is highly likely that photons measured from regions with many small/penetrating vessels have experienced multiple scattering from vessels with slow flow exhibiting MU_*n*=0.5_ dynamics and from vessels with higher flow exhibiting MO/SU_*n*=1_ dynamics. We hypothesized that these regions with mixed dynamics corresponded with the areas in which the static scattering and constant offset models did not improve, or improved less than expected, the model fits of the data (examples are shown with red ovals in Fig.3(C) and (F)). We consider two mixed dynamics models. The first considers mixtures of MU_*n*=0.5_ and MO/SU_*n*=1_ dynamics,

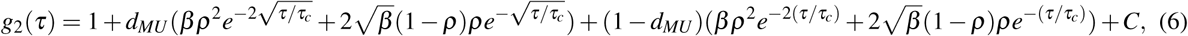

where the *d*_*MU*_ indicates the fraction of photons with dominant MU_*n*=0.5_ dynamics. We also consider the mixture of MO/SU_*n*=1_ and SO_*n*=2_ dynamics as

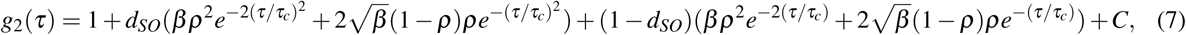

where *d*_*SO*_ indicates the fraction of photons with dominant SO_*n*=2_ dynamics.

The mixed dynamics model best fits are shown in Fig.3(G),(H),(I). From the F-test (Fig.3(I)), we clearly see that the mixed model does improve exactly the same regions that the other models failed to improve. This includes some parenchymal regions with a large number of penetrating vessels (red ovals in Fig.3(I)) as well as along the vessel walls of small surface vessels where shear induced diffusion of red blood cells introduces mixed dynamics^24^ (examples are shown with white ovals in Fig.3(I)). The best fit dynamic model shows that in these regions a transition between MU_*n*=0.5_ and MO/SU_*n*=1_ is observed, as indicated by the blue color (≈80% MU_*n*=0.5_, 20% MO/SU_*n*=1_) in Fig.3(G). Utilization of the mixed dynamics model also resulted in a reasonable estimate of the static scattering contribution in those regions, with 1 – *ρ≈* 0.3 instead of zero as observed for the static scattering model. The MO/SU_*n*=1_ to SO_*n*=2_ transition is not as evident, but does appear closer to the borders of large vessels, leading to the significantly better model fit in those areas.

#### Optimal model

While the simpler models are simplifications of the more complex mixed dynamics model, fitting more model parameters can increase uncertainty and introduce fit artifacts, such as a high amount of static scattering observed in the largest vessel where no static scattering is expected (compare Fig.3(E),(H)). We thus use the best model for each pixel as determined by the F-test using p<0.05 as the threshold for model selection. The results of the best model selection are shown in Fig.4.

**Figure 4.**
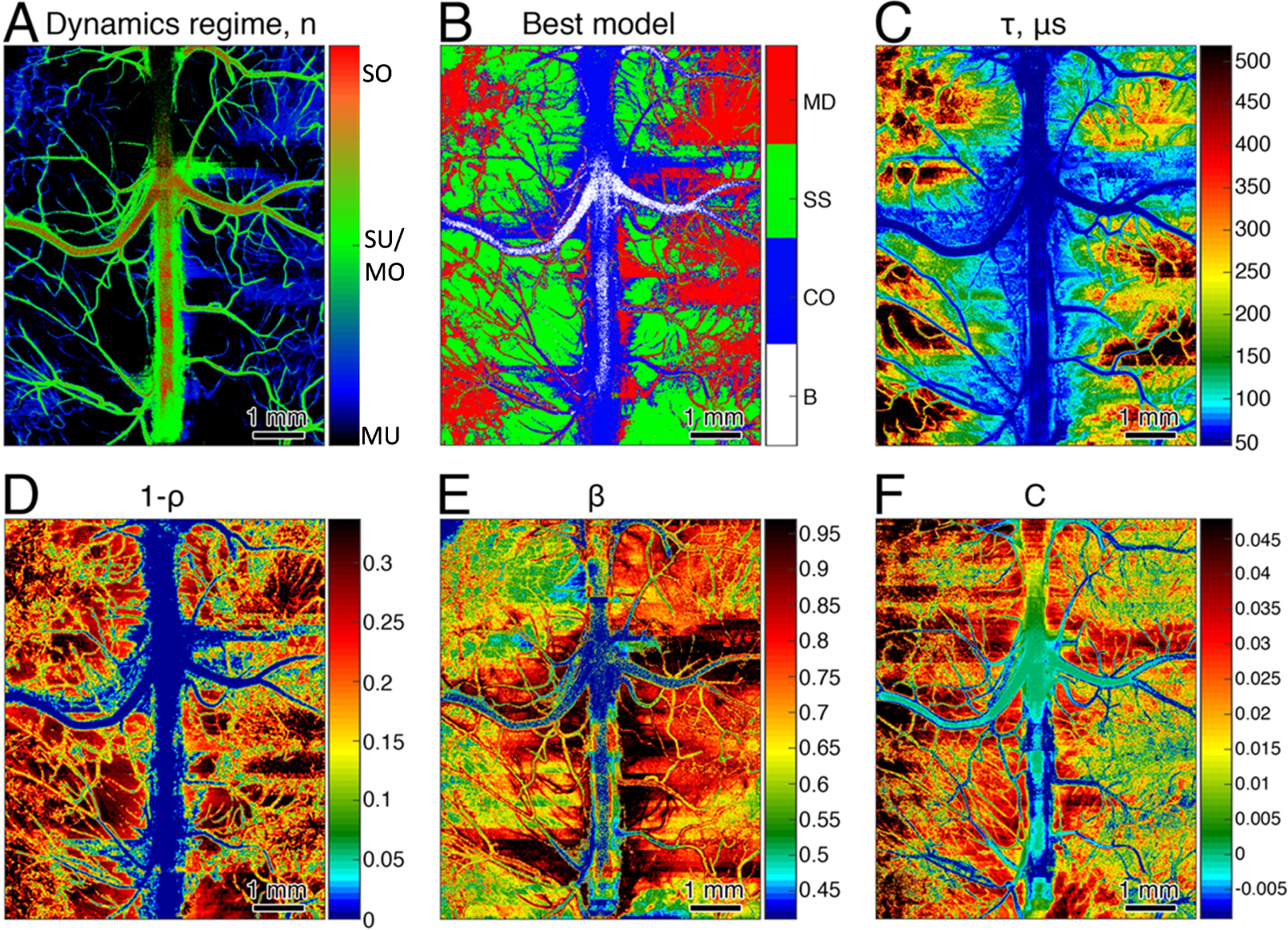
Best model selected based on the F-test. (A) - best model field correlation form. (B) - map of the best model selected according to the F test: Eq.1 (basic, notated as B on the colormap) - white, notated as B, Eq.4 (constant offset (CO)) - blue, Eq.5 (static scattering (SS)) - green, Eq.6,7 (mixed dynamics model (MD)) - red. (C)-(F) - corresponding parameters based on the best model selection.

The best fit model is shown in Fig.4(B). Following the F-tests from Fig.3, the largest vessels are best fit with the basic model (Eq.1). The constant offset model (Eq.4) provides the best fit for the immediate surroundings of the large vessel and the centers of medium sized surface vessels. The smallest surface vessels and parenchymal regions with a large number of penetrating vessels are best fit with the mixed model (Eq.6), while the rest of parenchyma is properly fit with the static scattering model (Eq.5). The contribution of static scattering is zero in large vessels and ≈0.25-0.3 everywhere else in the parenchyma. The estimate of the decorrelation time *τ*_*c*_ differs from that obtained with the basic model with *τ*_*c*_ ranging from 250 to 500 *μ*s (Fig.4(C)) instead of from 400 to 1500 *μ*s as observed with the basic model (Fig.2(B)).

### Stroke imaging - Improving Laser Speckle Contrast Analysis with Dynamic Laser Speckle Imaging

Knowledge of the dynamic light scattering regime depicted by the parameter *n* is critical for proper interpretation of blood flow and blood flow changes by dynamic light scattering and laser speckle contrast imaging methods. One of the major application areas for laser speckle contrast imaging and laser Doppler flowmetry for the past twenty years has been mapping and quantifying flow changes during ischemic stroke in animal models^27, 39–42^. Corrections to the laser speckle contrast theory^11, 14^ led to significant difference in the relative blood flow measurements made by LSCI today^18, 27, 42^ compared with the estimates from 15 years ago^40, 43^. Surprisingly, the corrected model resulted in LSCI, similar to LDF, measuring the relative blood flow in the ischemic core or occluded vessels to be above physiologically expected values^27, 32, 42, 44^, meaning it is likely that the relative flow change is not accurately quantified by LSCI, particularly for the larger flow changes expected in the ischemic core of a stroke. We hypothesized that one of the major reasons for this discrepancy, along with the static scattering effects^18^ which are also captured by DLSI, is the incorrect model of the field correlation function that has been used to analyze the data. In particular, it has been common to use the *n* = 1 form of *g*_1_(*τ*), but the analyses we presented above indicate that it is more accurate to use *n* = 0.5 for the parencymal regions. To test our hypothesis, we used both DLSI and LSCI approaches to measure the relative flow response (*rCBF*) during an MCAO induced stroke in a mouse model.

Results of the stroke experiment are shown in Fig.5. Relative cerebral blood flow (rCBF) calculated using the conventional LSCI model Eq.11 and Eq.12 results in rCBF=0.18 in the ROI associated with a penetrating arteriole affected by the occlusion, rCBF=0.33 in the parenchyma of the stroke core and rCBF=0.49 in the parenchyma in the immediate vicinity surrounding the core (Fig.5(A)). The respective *rCBF*_*DLSI*_ values, calculated using Eq.6 and Eq.13, are 0.06, 0.18 and 0.29 and align well with the physiologically expected results^42^. Use of the conventional 1*/K*^2^ LSCI model leads to an approximately 100% error in estimates of rCBF compared to the estimates by DLSI. Most of this error results from the conventional LSCI estimate of rCBF assuming *n* = 1. But even in the small penetrating arteriole where at baseline (i.e. pre-stroke), DLSI indicates that *n* = 1 is the proper model, the flow error is greater than 100%. This error occurred in the small arteriole because during the stroke DLSI indicates that the proper model is now *n* = 0.5. In other words, the flow reduction was sufficiently large in the small arteriole that the dynamic light scattering regime changed from *n* = 1 to *n* = 0.5. Furthermore, static scattering increased by ≈ 0.15 in the core of the stroke and by ≈ 0.07 in the immediate surroundings. The area of the flow drop and increase in static scattering is aligned with the area in the OCT angiogram image (Fig.5(E)) in which we no longer see capillaries or observe a reduction in their number. Ignoring such changes in the contribution of static scattering will result in errors in the estimate of the rCBF changes. In large vessel, which was not directly affected by the occlusion, the rCBF calculated by both methods is very close: *rCBF*_*LSCI*_ = 0.4 and 0.18 (ROI 4 and 5 correspondingly) and *rCBF*_*DLSI*_ = 0.36 an 0.17. This is to be expected since there is no change in the *g*_1_(*τ*) model nor in static scattering. Moreover it shows that the conventional LSCI model is not able to distinguish between large vessel with slow flow and occluded artery, as *rCBF*_*LSCI*_ is equal to 0.18 for both ROI 1 and ROI 5.

**Figure 5.**
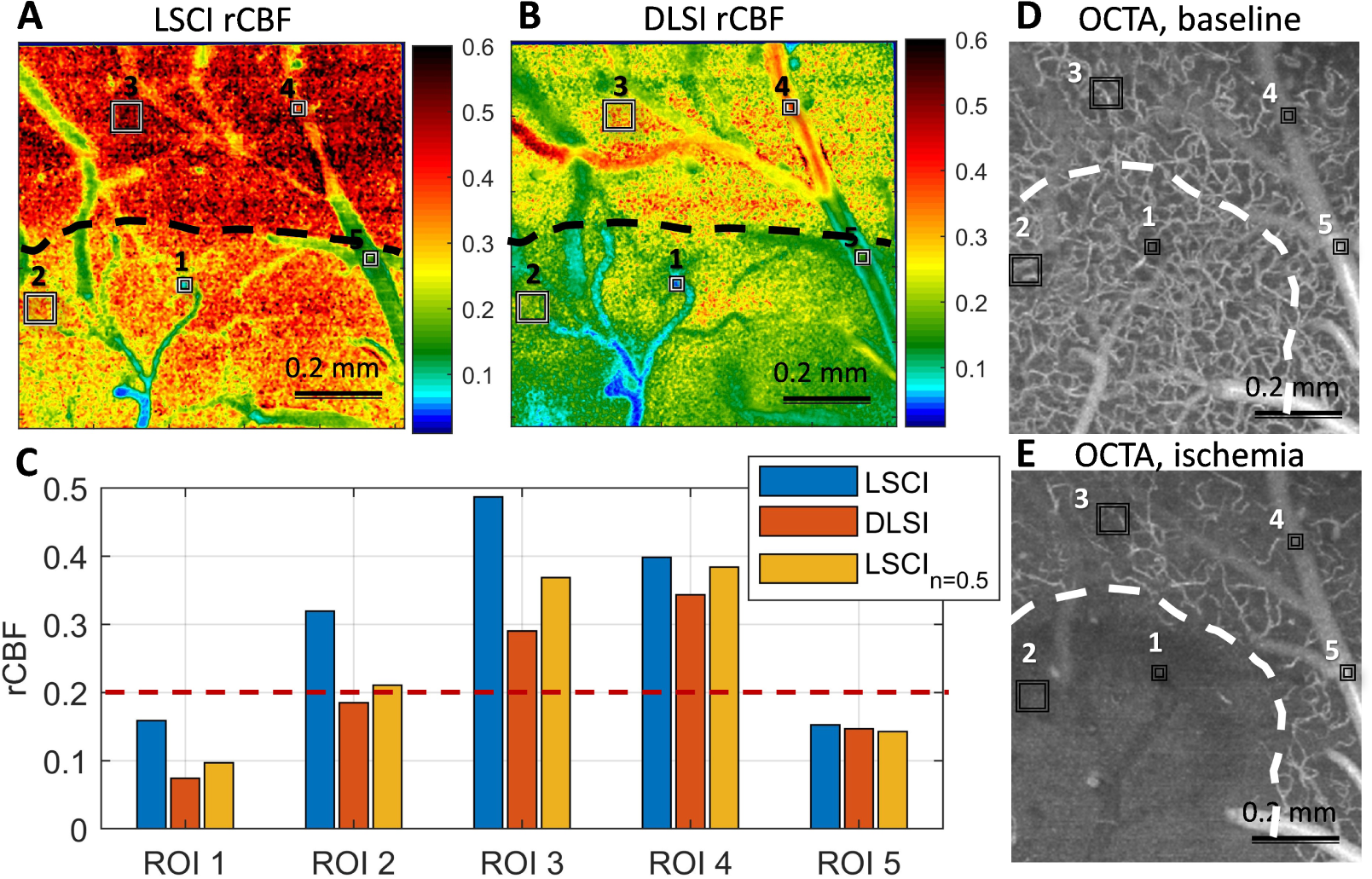
Application to the stroke imaging. (A),(B) - image relative cerebral blood flow after the MCA occlusion compared to the baseline flow. (A) - *rCBF*_*LSCI*_ (see Eq.12) calculated based on the spatial laser speckle contrast imaging using conventional 1*/K*^2^ model. (B) - *rCBF*_*DLSI*_ calculated based on the mixed dynamics model fit to the intensity correlation data. (C) - relative cerebral blood flow in selected regions shown in (A),(B), which correspond to occluded artery (ROI 1), parenchymal region in the core of the stroke (ROI 2), parenchymal region in the immediate vicinity of the core (ROI 3), regions of the large vessel which was not affected by the stroke directly (ROI 4 and ROI 5). Blue bars correspond to *rCBF*_*LSCI*_, orange to *rCBF*_*DLSI*_ and yellow to 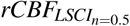, where the contrast is analyzed using Eq.9. Dashed red line shows the physiological stroke threshold of 20% of the baseline flow.One can see that *rCBF*_*LSCI*_ calculated using the conventional model drastically, by more ≈ 100% underestimates the flow change in the core of the stroke: with the flow in occluded artery and parenchyma being 0.18 and 0.33 compared to 0.06 and 0.18 measured with DLSI. Using contrast analyzed with Eq.9 significantly reduces the flow change underestimation. (D) and (E) - OCTA images of the corresponding region. Each image corresponds to the field of view of ≈ 1*x*1*mm*^2^. The core of the ischemic stroke is approximately indicated with white dashed line. The difference in the indicated area between LSCI/DLSI and OCTA is due to the later one scanning the sample at a slight angle.

**Figure 6.**
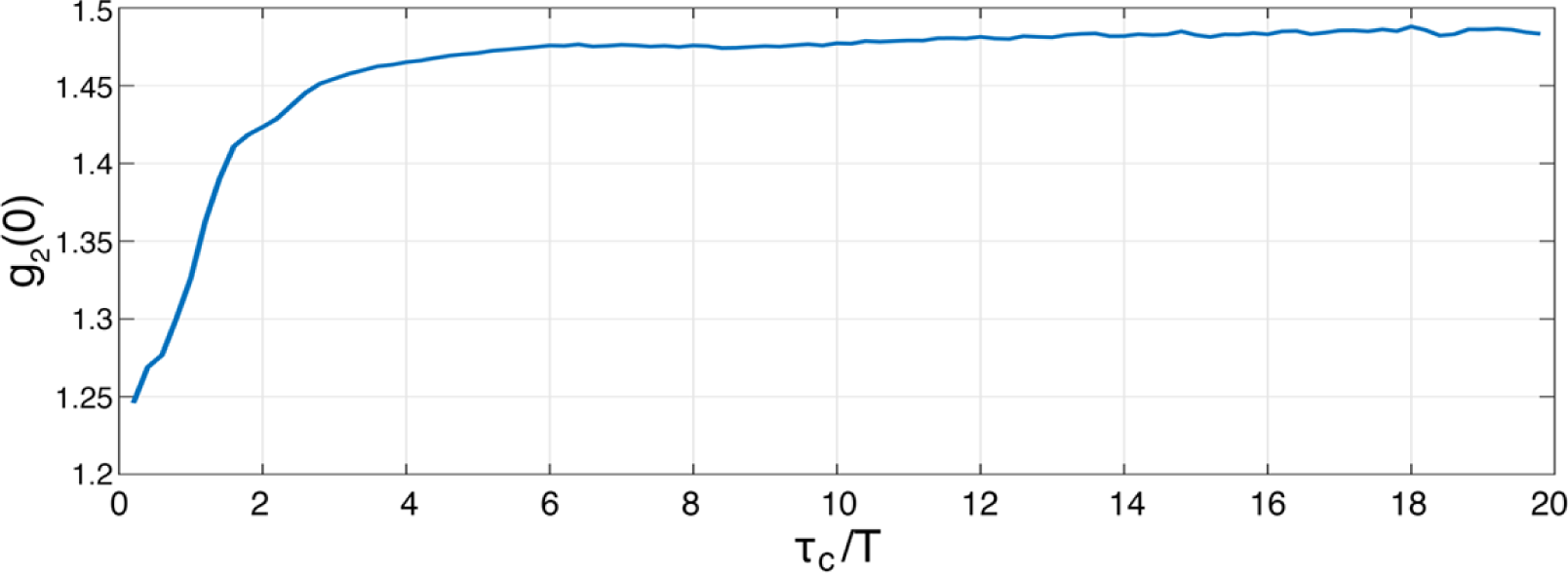
Supplementary, *g*_2_(0) dependency on the correlation to exposure time relation. *β*_*s*_ is not dependent on the *τ*_*c*_, and can be expected to be ≈0.47,defining the saturation value. *β*_*t*_, which depends on the correlation to exposure time relation is ≈0.5 for *τ*_*c*_*/T* = 1, ≈0.85 for *τ*_*c*_*/T* = 2 and is almost saturated after *τ*_*c*_*/T* = 4. Presence of static scattering leads to underestimation of the saturation value.

**Figure 7.**
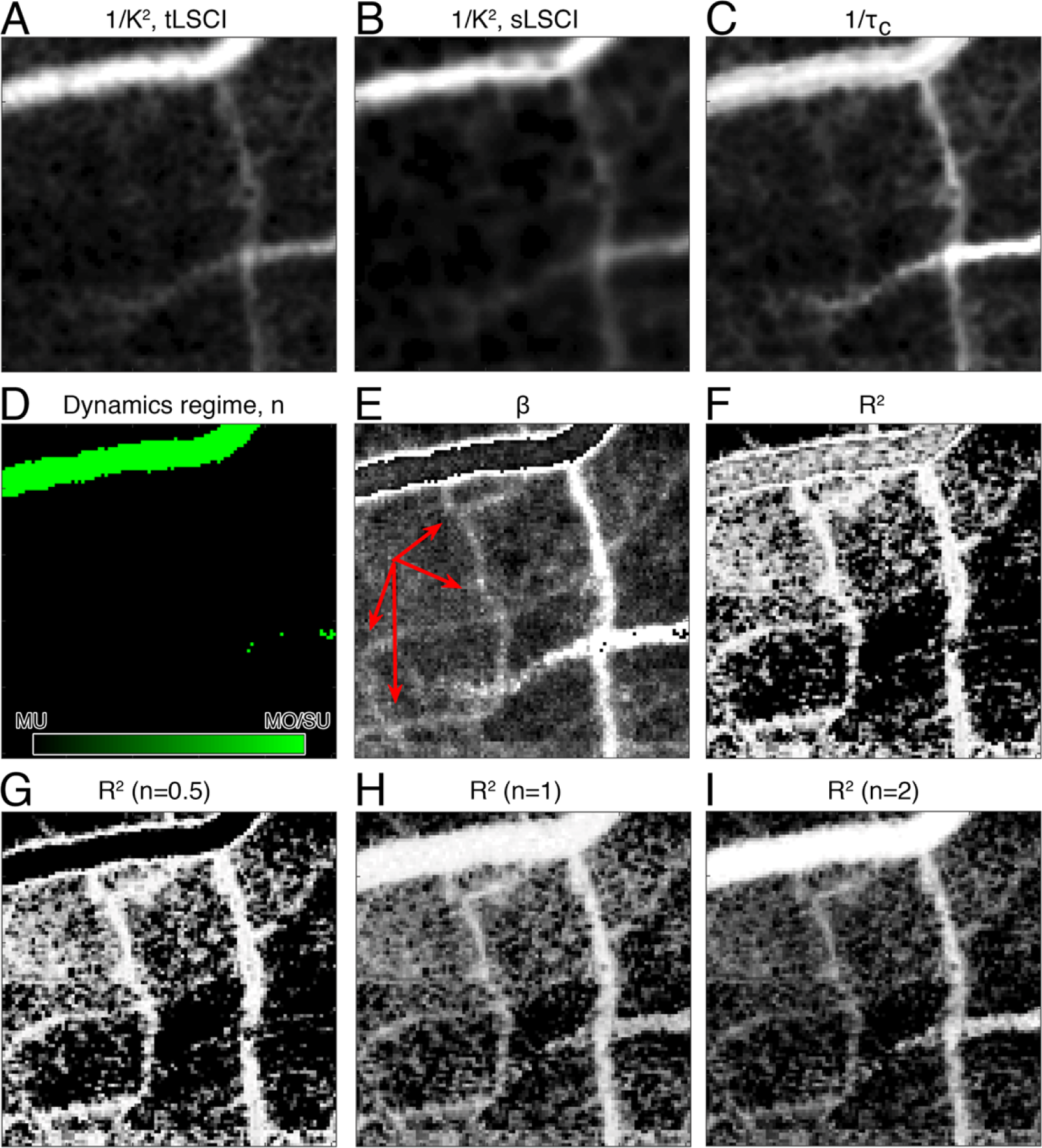
Supplementary, vessels visibility. A subset of 100 ×100 pixels of whole cortex image is shown. (A),(B) - flow visualization based on the temporal and spatial contrast analysis. (C) - flow visualization based on the decorrelation time constant *τ*_*c*_ obtained from the basic model fit. (D)-(I) - parameters and goodness of fit obtained from the basic model fit. It is clear that some vessels (examples are shown with red arrows in (E)) that are not visible in (A)-(B) and almost not visible in (C) appear clearly in panels (E)-(I).

**Figure 8.**
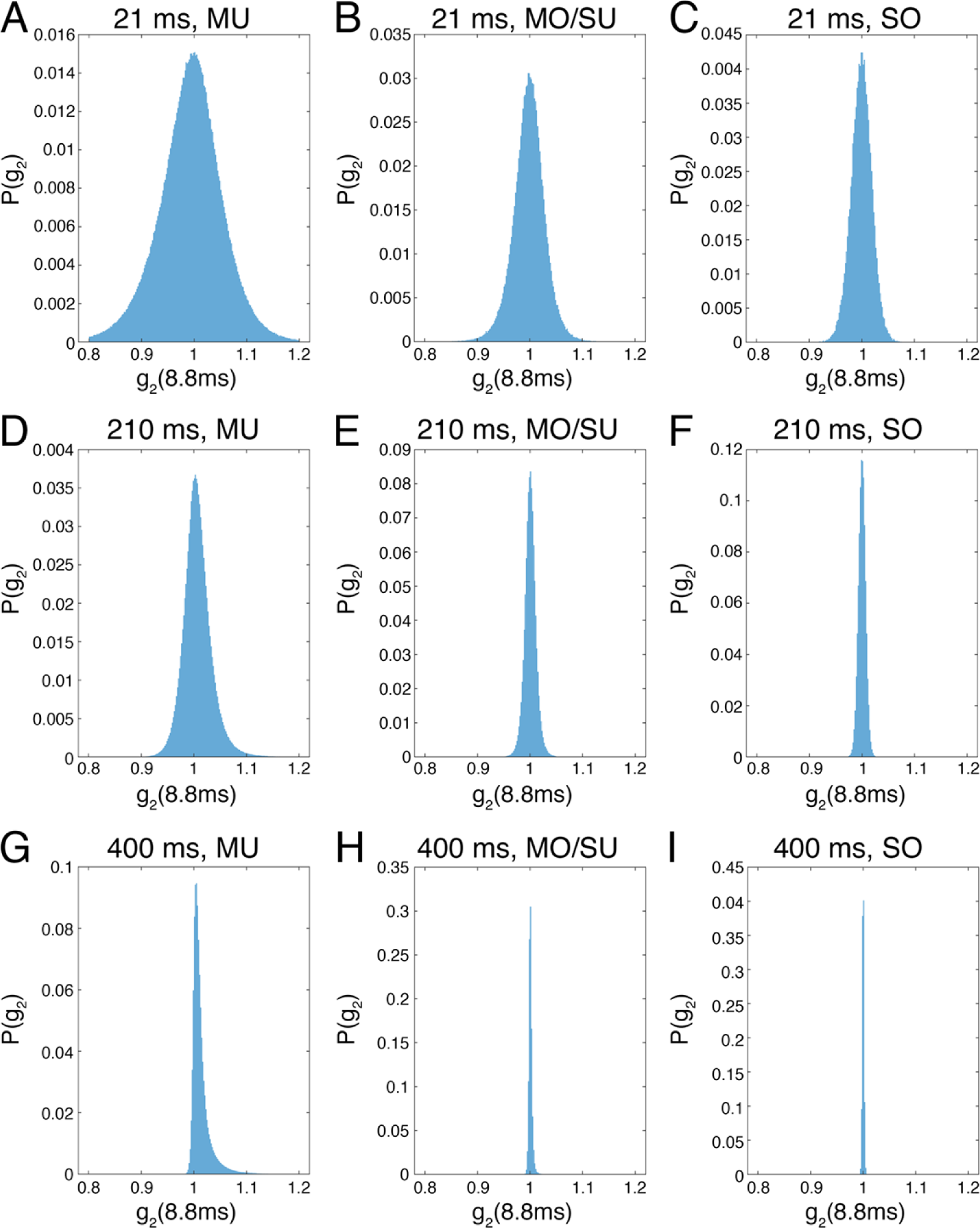
Supplementary, *g*_2_ distributions at the long time lag depending on the observation time. Columns correspond to pixels masked as MU (parenchyma), MO/SU (small and mid-sized vessels), SO (largest vessels) based on the best model fit. One can see the effect of statistical uncertainty from panels (A)-(F). It is pronounced stronger in pixels with longest decorrelation time (MU) and is reduced with longer observation times. In panels (G),(H),(I) which correspond to the 4s observation time the effect of statistical uncertainty is weak, but the positive component that reflects noise visibly shifts the mean to the value higher than one. The noise effect is particularly strong in the MU pixels, where it results in positive shift of the distribution peak to 1.005 value instead of 1.

**Figure 9.**
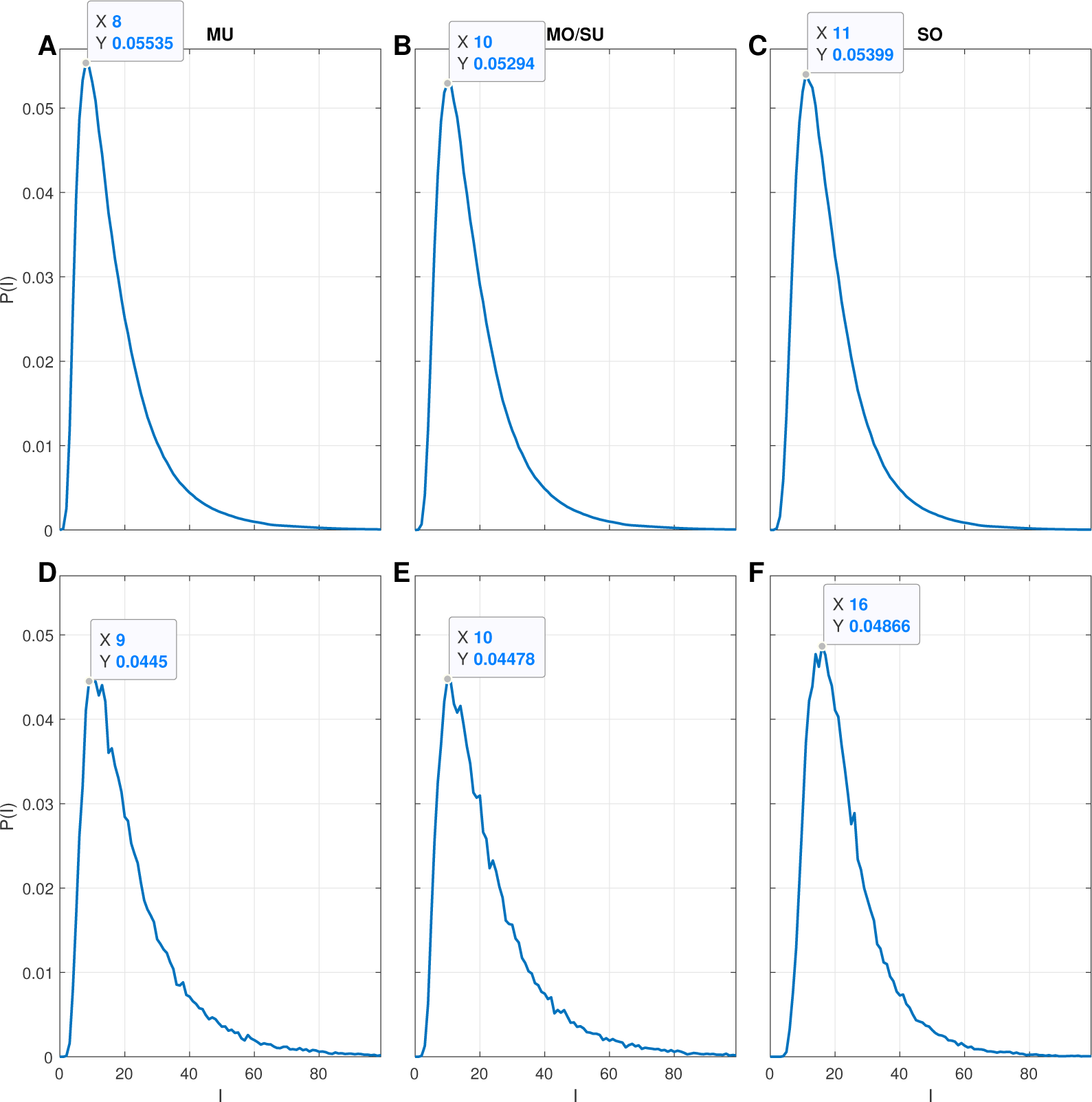
Supplementary, temporal intensity distributions at the long time lag depending on the observation time. Columns correspond to pixels masked as MU (parenchyma), MO/SU (small and mid-sized vessels), SO (largest vessels) based on the best model fit. (A),(B),(C) show temporal intensity distributions averaged over all of the pixels belonging to specific dynamics type, while (D),(E),(F) show exemplary distributions belonging to a single pixel of parenchyma, small surface vessel and large vessel respectively. The small shift of the intensity distributions which is observed for pixels with faster dynamics is caused by blurring of the speckle during the exposure time. This results in decreased beta in large vessels as well as makes surface vessels less sensitive to the camera noise.

**Figure 10.**
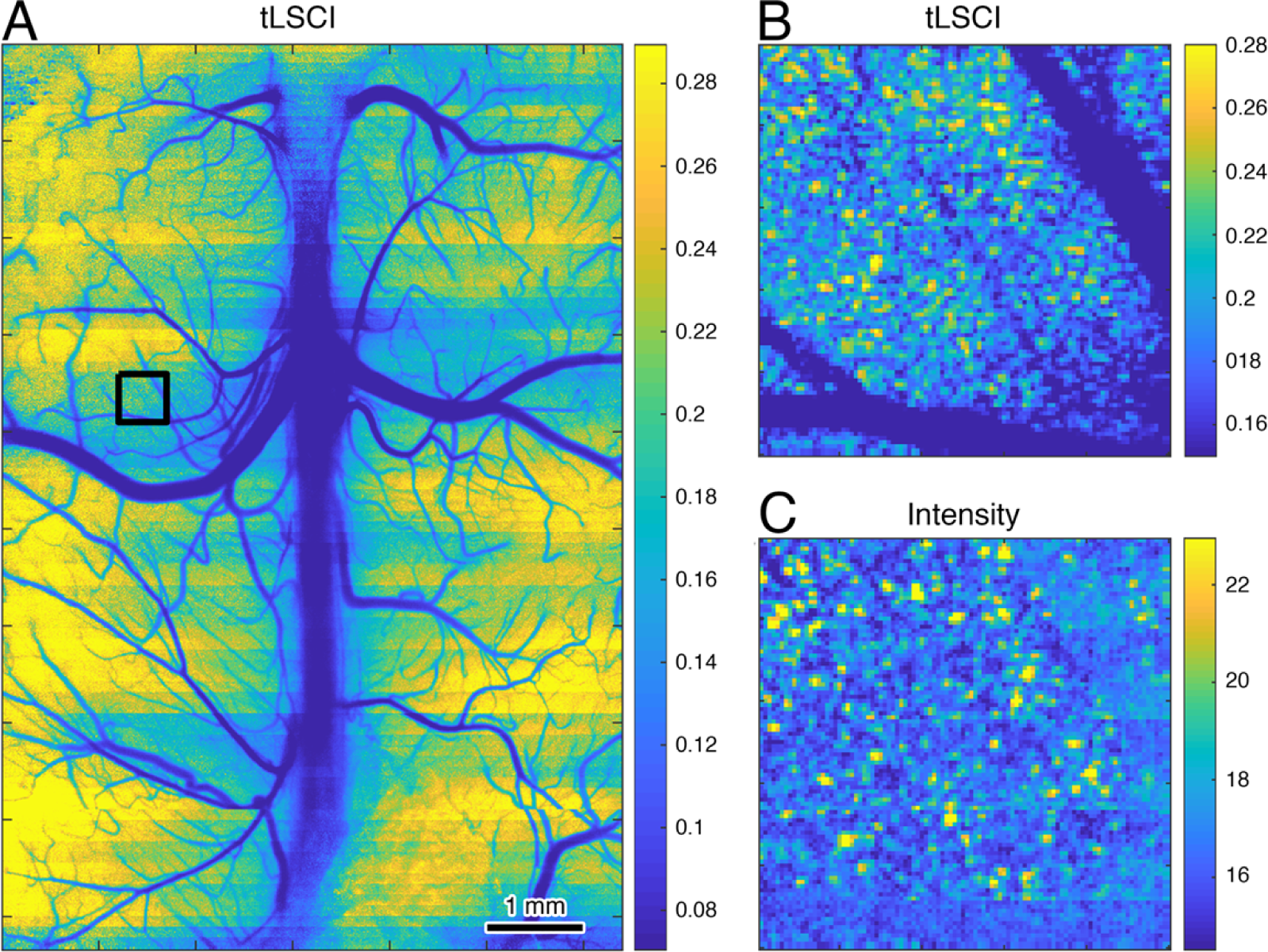
Supplementary, static scattering in temporal LSCI. (A) - tLSCI image calculated from the whole cortex data. (B) and (C) are contrast and intensity close-ups for the region marked in (A). One can see that because of the *I*_*c*_ ≠ ⟨*I*_*c*_ ⟩relatively weak static scattering appears as fluctuations of intensity and, consequently contrast. The fluctuation reach 20-50% and, contrary to the generally accepted opinion that tLSCI is less sensitive to static scattering, may lead to severe misinterpretation of the data.

**Figure 11.**
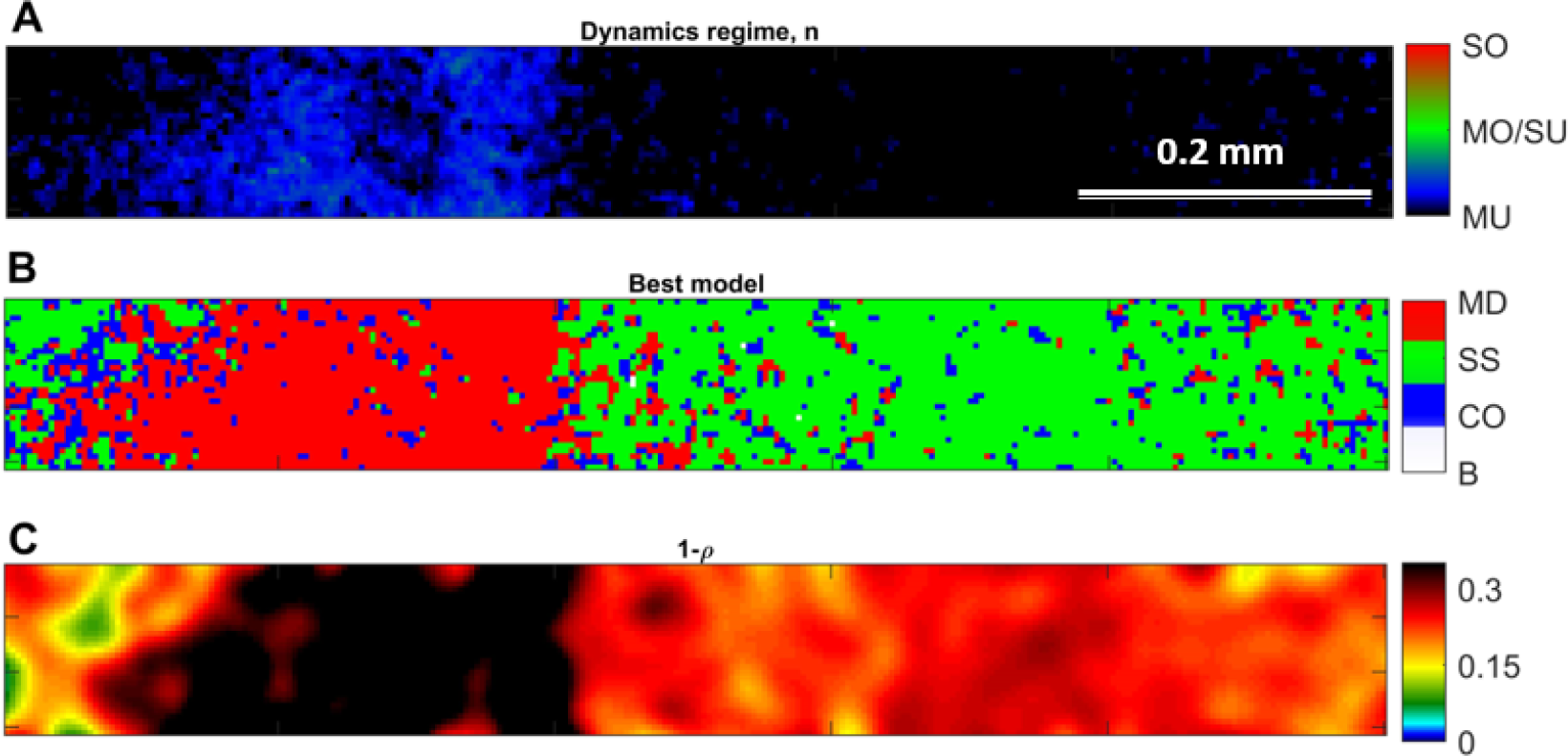
Supplementary, mouse skin imaging. (A) - best model field correlation form, (B) - map of the best model selected according to the F test, (C) - static scattering amount.

While this result shows that the use of the wrong *n* in the form of *g*_1_(*τ*) results in a significant error, it also guides us in improving the quantitative interpretation of the LSCI measurements. Following the theoretical derivation of the spatial laser speckle contrast theory in^11^ but using our mixed dynamics model for *g*_2_(*τ*), we have derived (see supplementary Eq.14-17) the relation for the MU_*n*=0.5_ to MO/SU_*n*=1_ transition as

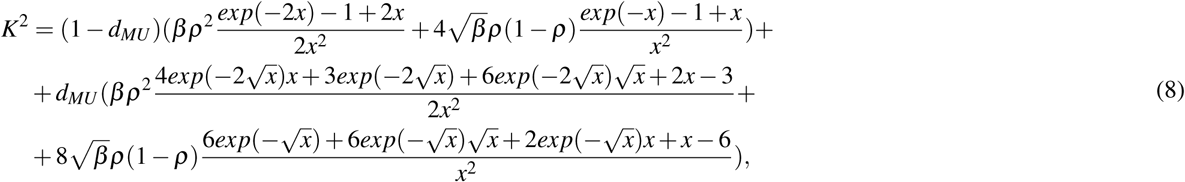

where *x* = *T/τ*_*c*_ and *T* is the camera frame exposure time.

For the parenchymal flow where we can assume *n* = 0.5, and further assuming that *ρ* = 1, *β* = 1 and *C* = 0, this simplifies to

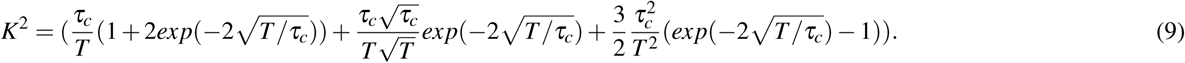

If we now reanalyze the LSCI data using the *n* = 0.5 model assuming *ρ* = 1 and *β* = 1, we then obtain rCBF of 0.10, 0.21 and 0.38 in the respective ROIs (see Fig.5(C)). These results are much better than when using the *n* = 1 model, but still have a ≈ 40% error compared to the DLSI results. We can also envision applications where data intensive DLSI could be used to calibrate LSCI using Eq.8 at select time intervals and then less data intensive LSCI is used to provide long term temporal sampling of rCBF changes.

## Discussion

We have introduced the Dynamic Laser Speckle Imaging technique, which allows us to measure the speckle intensity temporal auto-correlation function in a wide field and accurately estimate the correlation time and static scattering contribution using the most accurate model. We have shown how correlation decay varies in different regions of the brain, or, more specifically, in vessels of different sizes. The decay variation is regulated by the amount of photon scattering, the type and magnitude of motion exhibited by the light scattering particles, as well as by the presence of static scattering. In addition to the practical application of the technique for quantifying blood flow in biomedical applications, our results have answered longstanding questions associated with using laser speckle dynamics to measure blood flow, which will broadly impact studies utilizing laser speckle contrast imaging and laser Doppler blood flowmetry in diverse biomedical disciplines. In addition, our technique and analysis has raised new questions which will require further research.

### Dynamic scattering regimes

Arguably the most important and surprising result is the form of the field correlation function *g*_1_(*τ*), which is defined by the amount of photon scattering and type of motion and is represented by the value of *n* in Eq. 2. Historically, it was assumed that *n* = 1 in laser speckle contrast imaging^4, 9, 11, 12^ and multi-exposure speckle imaging^10, 18^. For laser Doppler flowmetry, values of *n* = 1 and *n* = 2 are both used without consensus on which is better^3, 6^. Our measurements of the temporal intensity correlation in the mouse brain, however, show that neither *n* = 1 nor *n* = 2 is correct per se, see Fig.4. The generally accepted *n* = 1 which reflects either single scattering with unordered motion or multiple scattering with ordered motion (MO/SU_*n*=1_) is present only in the “mid-sized” vessels, with diameters in the range of 20 – 200*μ*m. We observe that speckle dynamics in larger vessels demonstrate single scattering ordered motion behavior (SO_*n*=2_). And, contrary to expectations, we found that flow in the parenchyma, as well as in small visible vessels, display the behavior of multiple scattering with unordered motion (MU_*n*=0.5_). One can expect that it is not limited to the brain and *n* = 0.5 will be dominating speckle decorrelation for all tissues with a large number of small vessels and static scattering, such as skin (see Supplemental Fig. 11) or kidney. This result has a major effect on the accurate interpretation of LSCI, LDF and MESI measurements, as we saw in figure 5 when the wrong model was used with LSCI to interpret blood flow changes in a stroke model. Model selection error also likely explains the erroneously low flow changes reported in large retinal vessels during flickering light stimulation in^45^, in which SO_*n*=2_ dynamics are likely more appropriate while the paper utilized *n* = 1 dynamics.

While our results show that multiple scattering and unordered motion dominate the parenchymal blood flow, we are left with explaining the origin of the unordered motion dynamics. Interestingly, this observation of unordered motion of moving red blood cells has recently been explained for diffuse correlation spectroscopy, which is similar to DLSI in measuring the intensity temporal auto-correlation function, but does so with light that has typically diffused through a centimeter or more of tissue^7^. With DCS measurements of blood flow, unordered motion of red blood cells has also been observed, which has been explained by the shear-induced diffusion in vessels larger than ≈ 20 *μ*m in diameter^7^. This shear induced diffusion of red blood cells in these vessels has recently been confirmed microscopically with dynamic light scattering optical coherence tomography^24^. This, however, does not explain the DLSI observation of unordered motion in smaller vessels and the capillaries in the parenchyma, since one does not expect any shear-induced diffusion in these small vessels in which red blood cells experience plug flow and generally flow in a single file. Instead, we suggest that the unordered motion dynamics might be a result of red blood cell shape deformation which occurs in smaller vessels^46^. Additionally, the MU_*n*=0.5_ dynamics observed by DLSI in the capillaries of parenchyma might imply that DCS is more sensitive to the flow in small vessels than previously suggested^7^.

### Regime transition

Exploring the form of *g*_1_(*τ*), we showed that “mixed” forms were needed to best describe the data in some regions (Fig.4) of the mouse brain. Starting with the largest pial vessels, we observe in Fig.4(A) that SO_*n*=2_ dynamics occur in the center of the largest vessels that then transitions to MO/SU_*n*=1_ dynamics on the edges of these vessels which is likely single scattering from unordered motion of red blood cells resulting from shear induced diffusion of the red blood cells as observed by dynamic light scattering optical coherence tomography in^24^, rather than multiple light scattering from the ordered motion of red blood cells. Following these vessels to the smaller connected vessels, we see that the vessels are entirely depicted by MO/SU_*n*=1_ dynamics which is likely a result of increased shear throughout the smaller diameter vessels causing shear induced diffusion of red blood cells to dominate over the ordered laminar motion across the vessel diameter. Following along to the even smaller connected vessels, we now observe mixed MU_*n*=0.5_ to MO/SU_*n*=1_ dynamics. This is expected as the dynamics from the single scattering from the unorder motion in the smallest arterioles and venules is now slow enough that the contribution of the dynamics from the multiple scattering from the unordered motion of red blood cells in the capillaries becomes competitive. Modeling the MU_*n*=0.5_ to MO/SU_*n*=1_ transition proves to be important for obtaining reasonable fitting results with DLSI or MESI in parenchymal regions with “mixed dynamics”. From Fig.3(D),(E),(F) it is clear that the model with a pure rather than mixed form of *g*_1_(*τ*) fails to capture the static scattering component when *n* = 0.5, implying that 100% of the detected photons experienced dynamic scattering events. The reason is that the fitting algorithm compensates for the faster decay which is introduced by the partial presence of MO/SU dynamics by maximizing the amount of dynamic scattering, which, in turn, affects the estimate of *τ*_*c*_ leading to an error in the blood flow estimate. The issue is resolved by introducing the mixed dynamics and supported by the results of the F tests (Fig.4(F),(I))

### Static scattering imaging

DLSI permits estimation of the static scattering component. From Fig.4(D) we see that static scattering in the parenchyma ranges between 0.2 and 0.35, which corresponds well with previous estimations using MESI^18^. This means that ≈ 70% of the detected photons have experienced at least one dynamic scattering event. Given that no more than 4% of the brain cortex volume has moving red blood cells, the detected photons must be undergoing multiple scattering events in order of r ≈ 70% of them to experience a dynamic scattering event. Indeed, recent Monte Carlo simulations of photon migration through a microvascular network of the mouse cerebral cortex has indicated that on average the detected photons experience 15 to 20 scattering events^16^. This simply emphasizes that multiple scattering events should not be ignored.

### Insufficient Temporal Sampling

Two parameters in our mathematical description of the intensity auto-correlation function in DLSI require further discussion about their dependence on the camera exposure time *T*_*exposure*_ and the total observation time *T*_*total*_ (assuming no dead time between camera frames). These parameters are (i) the coherence parameter *β* which reflects source coherence and speckle averaging effects^11, 14, 21, 22^, and (ii) the constant term *C*, which reflects noise contributions^10, 11^.

Unlike LSCI or MESI, the DLSI equations do not explicitly include the exposure time *T*_*exposure*_, instead assuming that *τ*_*c*_ ≫ *T*_*exposure*_. While this assumption is true for most of the vessels, it breaks down for large vessels (> 100*μ*m) with faster flow, resulting in temporal averaging of the speckle dynamics during the measurement integration time. We have introduced this effect by denoting *β* as a *β*_*s*_*β*_*t*_ (*τ*_*c*_, *T*_*exposure*_) product, thus separating spatial and temporal averaging effects. The *g*_2_(0) dependence on the ratio *τ*_*c*_*/T*_*exposure*_ is shown in supplementary Fig.6. The value of *g*_2_(0) is a measure of *β* and since *β*_*s*_ is only dependent on spatial averaging effects, examining the dependence of *g*_2_(0) on *τ*_*c*_*/T*_*exposure*_ provides direct information about *β*_*t*_. The *β*_*t*_ value decreases for faster flow (i.e. shorter *tau*_*c*_) and represents reduced amplitude variation of the speckle intensity caused by temporal averaging. This makes it similar not only to the spatial averaging effects, but also to the contrast reduction of the laser speckle caused by the blood flow in LSCI. The *β*_*t*_ dependence on *τ*_*c*_*/T*_*exposure*_ can ultimately be modeled allowing one to exclude it from the fitting process.

In standard LSCI theory, the constant term *C* reflects the contribution from measurement noise^10, 11^. In DLSI, the constant term *C* also accounts for effects arising from an insufficient observation duration, *T*_*total*_, of the speckle dynamics. Given that the speckle fluctuations are occurring on a time scale of *τ*_*c*_, it is important that *T*_*total*_ ≫ *τ*_*c*_ to ensure that there is sufficient sampling of the temporal statistics of the speckle fluctuations. Generally, the standard deviation of the speckle intensity is equal to mean speckle intensity. But with insufficient temporal sampling, the standard deviation will not equal the mean. I<u>nstea</u>d, the proportionality will take on a value with a normal distribution that has a mean of 1 and a standard deviation of 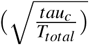. The effect of insufficient observation time is evident in supplementary Fig.8(A)-(F)). Even with long observation times, DLSI will still require the constant term *C* to account for camera readout noise and dark counts, just as is needed for LSCI. Theoretically, speckle intensity should be exponentially distributed, with speckles having zero or close to zero intensity appearing more often than speckles with higher intensity. In practice, however, camera read out and dark noise leads to a positive shift in the measured intensity distribution, resulting in < *I*(*t*)*I*(*t* + *τ*) > being greater than < *I*(*t*) >^2^ at large *τ* and thus *C* > 0.

### Temporal LSCI

As we discussed above following Eq.5, *I*_*c*_ *≠* ⟨*I*_*c*_ ⟩and *I≠* ⟨*I*⟩ for pixels with contributions from static scattering, leading to a pixel by pixel variation of the measured average intensity depending on the local static speckle contribution. This raises an important concern regarding the accuracy of relative blood flow changes using temporal laser speckle contrast analysis (tLSCI), which has been described as insensitive to the effects of static scattering^11, 30^. While a full mathematical derivation of spatial LSCI in the presence of static scattering exists^10, 11^, the theoretical derivation for temporal LSCI is lacking and it is simply conjectured that tLSCI can be used to get images with better spatial resolution because contrast is determined from temporal speckle fluctuations for each pixel and not spatial speckle fluctuations across a neighborhood of pixels^11, 30^, even in the presence of static scattering. The problem is that contrast is given by the standard deviation divided by the average intensity, and although the standard deviation is accurately estimated from pixel temporal fluctuations, the average intensity will have pixel by pixel variation due to the presence of static scattering. This static speckle pattern will lead to variations of contrast for the pixels with the same *τ*_*c*_ and < *I*_*f*_ > as one can see in supplementary Fig.10. The consequence is important; tLSCI, in general, should only be used for the studying ergodic systems in which a temporal average equals the spatial average. Given that parenchymal regions of the brain have significant contributions from static scattering, even when no skull or dura matter is present, tLSCI cannot be used to get accurate measurements of blood flow without any spatial averaging.

### Improving LSCI analysis

We have demonstrated that DLSI can be used to guide model selection to improve the analysis of the laser speckle contrast data. For the parenchymal flow, a simplified contrast equation (Eq.9) can be used to drastically improve the accuracy of relative cerebral blood flow changes measured with LSCI. This is particularly crucial in conditions where large flow changes are expected, such as in ischemic stroke. As we show in Fig.5(C), assuming the *MU*_*n*=0.5_ model with *β* = 1 and *ρ* = 1, we obtain relative flow which is much closer to the DLSI measurements and corresponds to physiologically expected values much better than the values obtained with the original LSCI analysis based on the *MO/SU*_*n*=1_ model. This result emphasizes the importance of using DLSI to determine the correct form of the dynamic model (i.e. *n*) to use to accurately analyze each pixel in the image; not just for brain but for the application of laser speckle analysis of blood flow in all organs. For instance for skin one can expect *n* = 0.5 or mixed dynamics everywhere (see Supplemental Fig.11), meaning that current applications of LSCI for imaging the skin are underestimating flow changes. It should be expected that clinical applications such as diagnosis of skin burns^47^ will be strongly affected, as well applications imaging the kidney^33, 48^ and liver^49^. For imaging large retinal vessels, single scattering and ordered motion (i.e. *n* = 2) might dominate instead, which will affect the interpretation of data acquired by such tools as LSFG NAVI^45^. One can easily benefit from knowing the appropriate field correlation model by either recalculating the flow change depending on the location of the region of interest, or by using a digital mask that separates tissue perfused mostly with capillaries from larger vessels and calculating the flow change using the appropriate model for each region.

Furthermore the interpretation of LSCI can be further improved by using parameters derived from the DLSI fit and using Eq.8 to analyze the contrast. Only a few seconds of DLSI data acquisition is required to get the proper form for *g*_1_(*τ*) and estimates for *β* and *ρ*, which can be used with to convert LSCI contrast values to the decorrelation time *τ*_*c*_ (and thus blood flow index) with improved accuracy. While for stroke imaging it makes more sense to simply use DLSI estimates to get the flow and static scattering variation, we do envision other applications where the data and computationally intensive DLSI can be used during the baseline to calibrate the less data and computationally intensive LSCI. This may become particularly valuable for applications where no change in static scattering is expected, allowing the DLSI calibration to transform LSCI into a quantitatively accurate tool providing quantitatively accurate estimates of blood flow with high temporal resolution.

## Methods

### Animal preparation

All animal procedures were approved by the Boston University Institutional Animal Care and Use Committee and were conducted following the Guide for the Care and Use of Laboratory Animals. Two experiments were designed: one for the imaging of the whole cortex and another one for monitoring of the blood flow change during the stroke.

#### Whole cortex imaging

C57Bl6 mice were anesthetized with isoflurane (3% induction, 1–1.5% maintenance, in 1L/min oxygen) during surgery and imaging sessions. After removal of the scalp, the skull was removed to fit the placement of the Crystal Skull^50^ The glass then was sealed with dental acrylic and the animal was recovered for 3 weeks before the imaging session. During surgery and imaging, heart rate and oxygen saturation was non-invasively monitored (Mouse Stat Jr, Kent Scientific) and all noted measurements were within the expected physiological range.

#### Stroke imaging

A 15-week-old male C57BL/6 mouse was used for the stroke experiment. The animal was anesthetized with isoflurane (2-3% induction, 1-2% maintenance, in 60% Nitrogen, 40% Oxygen mixture. Following a midline skin incision, and removal of scalp and periosteum, a custom-made aluminum bar was glued over the left half of the skull, for fixation of the head. Then, approaching between the right lateral epicanthus and external auditory meatus, the temporalis muscle was dissected to reveal the squamous portion of the temporal bone. The temporal bone over the distal middle cerebral artery (MCA), 1 mm above the zygomatic arch was drilled and a craniotomy of 2 mm diameter was opened, with dura intact. To prevent overheating, extended drilling of the same area for more than 2-3 seconds was avoided and the skull was cooled with aCSF at room temperature throughout the drilling process. The exposed MCA and surrounding cortex were flushed with warm (at 37 C*°*) artificial CSF (aCSF) and then was covered with aCSF-soaked-gauze for protection. Next, another frontoparietal craniotomy was opened (4 mm in diameter) to visualize the MCA-supplied cortex. The cortex was then covered with 0.7% agarose solution in aCSF, followed by a 5-mm glass coverslip. The window was sealed with dental cement. The animal was then placed under the imaging system. We waited for at least an hour for stabilization of the cerebral blood flow until we started the baseline imaging. During the entire length of the surgical procedure, the animal was heated by a homeothermic blanket with rectal-probe feedback to maintain the temperature at 37 C*°*. Mean arterial pressure was monitored in animals with femoral cannulation. Arterial oxygen saturation and heart rate were monitored noninvasively with a paw probe (MouseSTAT Jr., Kent Scientific Instruments).

We used a ferric-chloride induced middle cerebral artery thrombosis model, as previously described^42^. After the acquisition of baseline imaging data, a piece of filter paper (0.3×0.5 mm, Whatman No.1) soaked in 30% ferric chloride (FeCl3, the solution in isotonic saline) was placed over the intact dura matter along the trace of the MCA right after the zygomatic arch. This concentration of FeCl3 was preferred to prevent spontaneous recanalization of MCA, based on previous experience^42^. The paper was kept in place for 10 minutes, then the flow in pial MCA branches was checked with laser speckle contrast imaging. Since residual flow was detected, another piece of soaked paper was reapplied for 5 minutes and repeated laser speckle contrast imaging confirmed dMCA occlusion. The FeCl3 soaked-paper was then removed and the dMCA trunk area was washed with warm aCSF.

### DLSI

A high-speed camera (1280×1024 pixels, 991 fps, 5 *μ*m pixel size, Fastec IL5-S, USA) was used to record the backscattered light, through a 5x (whole cortex imaging) objective with NA= 0.14 or a 10X (stroke imaging) objective with NA= 0.28 (Mitutoyo, Japan). A polarizer was placed in front of the objective to increase the contrast range. The number of active pixels was reduced to an image stripe of 1280×32 and the exposure time *T*_*exposure*_ was set to 31*μ*s, allowing the camera to reach the maximum frame rate of 22,881 frames per second. Coherent light was delivered to the object using a free space volume holographic grating (VHG) stabilized laser diode^21^ (785nm, LD785-SEV300, Tholrabs, USA) operated at the recommended settings. The light was collimated, passed through the isolator (IO-5-780-VLP, Thorlabs USA), expanded along the long axis of the camera frame with an anamorphic prism pair (PS875-B, Thorlabs, USA) and further expanded by a beam expander (GBE10-B). This resulted in an ≈ 10×2 mm, 200mW power beam spot at the surface of the brain which was then aligned with the camera’s field of view as shown in the bottom right part of Fig.1(A). The size of the speckle on the camera was adjusted by altering the pupil diameter of an iris in the detection path to achieve a speckle to pixel size ratio of approximately 2. To scan the whole field of view when imaging the whole cortex (an approximate 8×6 mm imaging area), the stage with the animal (LTS150, Thorlabs, USA) was translated in the Y direction such that sequential image stripes had an overlap of 10 pixels to permit registration of sequential stripes. For imaging the smaller field of view in the stroke experiment, the approximate 1×1 mm imaging area was contained within the illumination spot and thus we did not need to translate the sample but instead translated the 1280×32 image stripe on camera sensor. Each 1280×32 stripe was recorded for 4 seconds, resulting in 91,544 speckle images per stripe.

### Data analysis

#### Intensity correlation

The image stripes were stitched together by spatial co-registration of the *τ*_*c*_ images and the overlap was removed. The camera black level was evaluated and subtracted from the intensity prior to the analysis. The resulting speckle images were processed in order to obtain the speckle intensity temporal autocorrelation function *g*_2_(*τ*)^35–37, 51^:

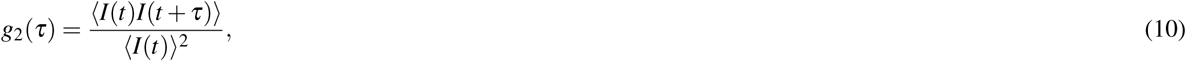

where *I* is the intensity recorded at a specific pixel, *t* is a time corresponding to the current frame and *τ* is the time lag value. Angle brackets denote averaging across the observation time *T*_*total*_ = 4s. Depending on the experiment, the time lags *τ* ranged from either 0 to 8.8ms for whole cortex imaging or 0 to 17.2ms for the stroke imaging. Longer time lags were used for stroke imaging because of the slower blood flow and thus slower decorrelation. **Model fitting.** The calculated intensity correlation was fitted with the models indicated in Eq.1,4-7 using a nonlinear least squares fitting algorithm in Matlab 2018b with the following parameters: the minimum change in the coefficients for finite difference gradients was 10^*-*12^, the maximum number of evaluations of the model was 3000, the maximum number of iterations allowed for the fit was 1500, the termination tolerance on the model value was 10^*-*12^, the termination tolerance on the coefficient values was 10^*-*12^. All other parameters were set to the default Matlab 2018b fit options. The fitting parameters were constrained to 0..1 for *β, ρ, d*_*MU*_, *d*_*SO*_; −1..1 for *C* and 0..8.8ms or 0..17.2ms for *τ*_*c*_ for whole cortex and stroke imaging respectively. Initial conditions for the decorrelation time constant *τ*_*c*_ were identified by finding the time lag at which *g*_2_(*τ*) crosses the value 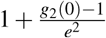^52^. The value *g*2(0) was excluded from the fitting data for all pixels, due to the uncorrelated acquisition noise strongly affecting the zero lag correlation. The “tail” of the intensity correlation curve, starting from the point where the correlation drops below 10% of the initial value was also excluded from the fitting data, except for the cases where the total number of points was less then 5 (large vessels with fast decorrelation). In those cases, 5 points, starting from the first lag were used. **Best model selection and F test**. When comparing the models with the same number of parameters, for example when the best fit dynamics regime (*n*) is identified for the specific *g*_2_(*τ*) model (Fig.2(A), Fig.3(A),(D),(G)), the best model was chosen as the model with the largest *R*^2^ value. When comparing models with a different number of parameters the F test^38^ was performed (Fig.3(C),(F),(I)). The model with more parameters was considered to be a better model when *p – value* ≤ 0.05 (see Fig.4(B)).

#### Contrast analysis

To obtain contrast images from the same data set - sequences of 114 speckle images were averaged to providing a close equivalent to the image captured with a 5ms exposure time. The contrast was calculated and converted to the blood flow index (BFI) according to the commonly used simplified model^26–29, 53^

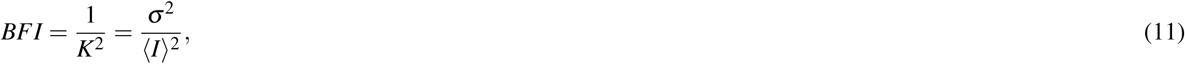

where *K* is contrast, *σ* and ⟨ *I* ⟩ are the intensity standard deviation and mean of the pixel values over either a 5×5 neighborhood (spatial contrast^11^) or 25 frames (temporal contrast^30^). This resulted in 803 (spatial contrast) or 32 (temporal contrast) images that were then averaged, providing the contrast information averaged over the entire observation period *T*_*total*_ = 4s. **Relative blood flow**. Relative blood flow changes during stroke was calculated from the contrast data as:

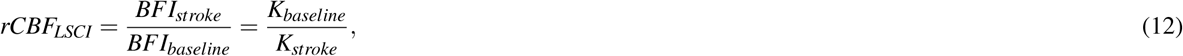

where *K*_*baseline*_ and *K*_*stroke*_ are the spatial contrast calculated according to Eq.11 during the baseline and ischemic stroke respectively. The relative blood flow changes measured with DLSI were calculated as:

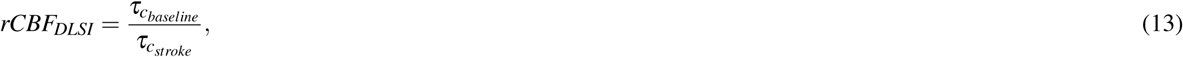

where 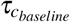 and 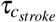 are the correlation times obtained by fitting the chosen model to the corresponding experimental data. The code for the intensity correlation and contrast calculations was written in Matlab and executed using the GPU to achieve a reasonable performance. The code is available in the BU Neurophotonics GitHub repository^54^.

### OCTA

For the stroke experiment, optical coherence tomography angiography was performed. A spectral-domain OCT system (1310 nm center wavelength, bandwidth 170 nm, Thorlabs Telesto III) was used for imaging of the cerebral cortex. The axial resolution of the system in the air was 4.6 mm. Given the light refractive index of 1.35 for brain tissue, the axial resolution in the brain was 3.5 mm. The imaging speed was 76,000 A-scan/s. A 10X objective (10X Mitutoyo Plan Apochromat Objective, NA) was used in this study, providing a transverse resolution of 3.3 mm. The micro-vascular angiogram was constructed by a decorrelation-based analysis method of the OCT data^55^. OCT data was acquired over an area spanning 500×500 pixels equivalent to 1000×1000×1000 *μ*m3. Each angiogram volume acquisition took ≈9 seconds. Ten volumes were acquired and then averaged to increase the signal-to-noise ratio. For the image present, the maximum intensity projection (MIP) over a depth range of 100-200 *mu*m beneath the brain surface were calculated from the angiogram volumetric data.

## Acknowledgements (not compulsory)

D.D. Postnov was supported by grant NNF17OC0025224 awarded by Novo Nordisk Foundation, Denmark. Support was also provided by NIH R01-MH111359, R01-EB021018, and R01-NS108472.

## Author contributions statement

DDP and DAB conceived the model; DDP, DAB and ESE conceived the experiments; DDP, ESE and KK conducted the experiments, DDP, ESE and JT analysed the results. All authors reviewed the manuscript.

## Supplementary data

### Derivation of Eq.8 and Eq.9

Second moment calculation^11, 14^:

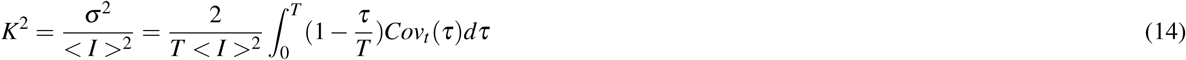

Temporal autocorrelation relation to covariance^11, 14^:

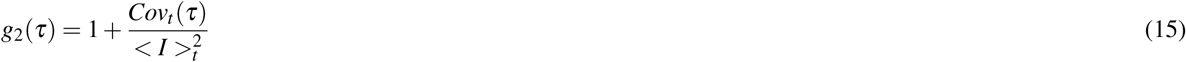

Substitute *g*_2_(*τ*) in Eq.15 with Eq.6:

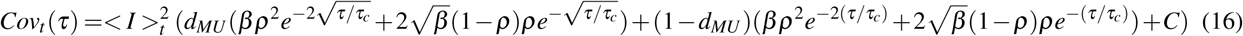

Substitute *Cov*_*t*_ (*τ*) in Eq.14 with Eq.16 (detailed integration steps are omitted):

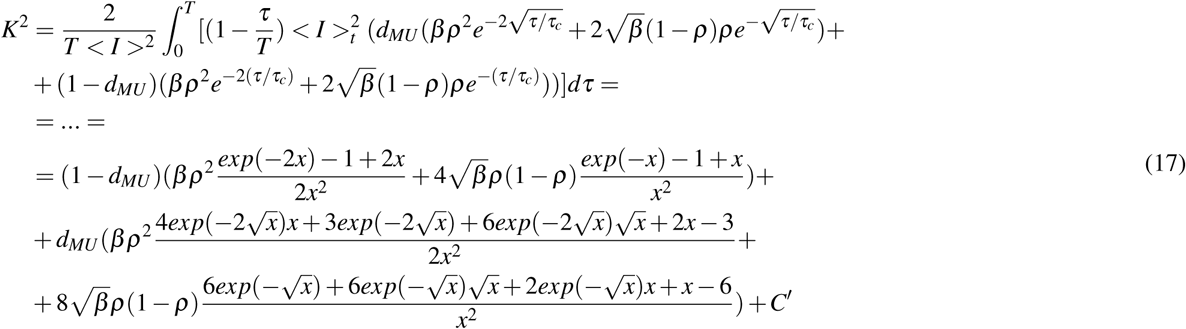

Assume *C*’ = 0 to obtain Eq.8 and *C*’ = 0,*d*_*MU*_ = 1,*ρ* = 1 and *β* = 1 to obtain Eq.9.

